# Investigating the Impact of Offer Frame Manipulations On Responders Playing The Ultimatum Game

**DOI:** 10.1101/2021.07.06.451289

**Authors:** Eve Florianne Fabre, Rino Rumiati, Cristina Cacciari, Sylvie Borau, Mickael Causse, Lorella Lotto

## Abstract

The present study was designed to test the impact of frame manipulations on the decision-making of responders playing the ultimatum game. Experiment 1 investigated responders’ event-related potentials (ERPs) measured in response to the offers as a function of the frame (i.e., negative: “the proposer *keeps”* versus *positive “the proposer offers”*). While no difference in acceptation rate was found as a function of the offer’s frame, electrophysiological results suggest that the stronger negative affective response to the offers in the negative frame (N400) was successfully reappraised by the responders (P600), possibly explaining why the offer frame manipulation did not modulate acceptation rates. No framing effect was found when the ultimatum game was played in its one-shot version (Experiment 2), suggesting that repeated measurements did not affect responders’ behavior. However, an offer framing effect was found in female (but not in male) responders, when the players’ cognitive charge was increased using more complex game rules (Experiment 3), presumably reflecting women’s greater affective responses to negative outcomes. Taken together, these results suggest that framing manipulations are associated with complex affective and cognitive processes, supporting the cognitive–affective tradeoff model.

## 1. INTRODUCTION

The framing effect occurs when equivalent descriptions of a decision problem lead to systematically different decisions (Tversky, & Kahneman, 1981). This phenomenon has been widely investigated over the last three decades and was shown to affect decisions in many fields, such as medicine (e.g., Gong, et al., 2013), politics (e.g., Aalberg, et al., 2011) or economics (e.g., De Martino, et al., 2006) among others.

### 1.1. Theoretical approach of framing effects

Normative models and more specifically game theory (von Neumann & Morgenstern, 1947) and its fundamental axiom of “extensionality” (Arrow, 1982), also called “invariance” axiom (Tversky & Kahneman, 1986), posit that the manner the different options of a decision problem are presented should not affect the decision-making of individuals. Allais (“Allais Paradox”, 1953; Allais & Hagen, 1979) followed by Kahneman and Tversky (1979) were the firsts to challenge the invariance axiom. By demonstrating the existence of framing effects, i.e., a deviation from rational decision-making, these authors attested the limitations of most decision-making normative models assuming description invariance (Fox & Poldrack, 2008; Tversky, & Kahneman, 1981). At variance with the latter models, the framework of prospect theory explicitly acknowledges the fact that choices are dependent on the way the different options of a problem are described (Kahneman & Tversky, 1979). This theory posits that (1) people analyze the different options they have, in terms of potential gains or losses compared with a neutral reference point, (2) are more sensitive to losses than to corresponding gains, and (3) tend to make more risky decisions in the loss domain than in the gain domain. Later in time, various other theories were proposed to explain the occurrence of framing effects (Gonzalez, et al., 2005). First, affective theories consider the difference in emotional value attributed to the alternatives of a problem depending on its frame (e.g., Mellers, et al., 1999) and posit that losses loom larger than equivalent gains because individuals assign more value to feelings of displeasure than to feelings of pleasure. Second, cognitive theories posit that the cognitive effort necessary for the valuation of alternatives may vary according to the frame used to describe these alternatives and that individuals tend to choose the easiest apparent correct decision they can find between the alternatives (Payne et al., 1993). Finally, the cognitive–affective tradeoff model brings cognitive and affective theories together and posits that depending on the frame, the alternatives may be associated with different cognitive costs and affective values, triggering a variation in the preference for an alternative or another (Gonzalez et al., 2005; Kahneman & Frederick, 2007).

The impact of framing effect on economic decision making has been largely investigated using a wide range of experimental paradigms (e.g., De Martino et al., 2006; Guo et al., 2013; Windmann, et al., 2006; Zhou & Wu, 2011), including the ultimatum game (Sarlo et al., 2013; Tomasino et al., 2013). As the ultimatum game is one of the main experimental paradigms used in the investigation of fairness-related decision making (Camerer, 2003; Sanfey, 2009), determining whether the way the offers are framed in this economic game modulates the responders’ decision-making is of great importance to prevent potential experimental biases. Before presenting the aim of the present study, we will first present the ultimatum game paradigm and the main studies investigating the responders’ behavior and the previous studies that investigated the impact of framing manipulation on responders’ decisions.

### 1.2. The ultimatum game

In the ultimatum game, a proposer is given a fixed amount of money (e.g., 10 euro) s/he has to split with a responder (Güth, et al. 1982). If the responder accepts the offer, both players get their shares as proposed by the proposer whereas, if the responder decides to reject the offer s/he has been made, none of the players gets the money. The expected utility theory predicts that aiming at maximizing their outcomes, the proposer and the responder should respectively offer the smallest share possible and accept every proposed share without distinction (von Neumann & Morgenstern, 1947). However, many studies have shown that the great majority of people deviate from utility-based expected behaviors (Camerer, 2003). Proposers typically offer shares close to a 50-50 division (Thaler, 1988; Roth, 1995; Güth, 1995; Camerer & Thaler, 1995) and responders very often reject low shares (i.e., 20% of the total or less; Güth et al., 1982; Camerer, 1999, 2003; Sanfey, 2009).

Responders were found to adopt ready-to-go responses based on internal heuristics (i.e., an easy to implement short cut “rules of thumb”), consisting of accepting fair shares and rejecting unfair ones quite automatically (Civai, 2013). The automatic rejection of unfair shares results from the negative affective response (e.g., anger, disappointment, frustration) triggered by the reception of a share deviating from equity (Pillutla & Murnighan, 1996; Sanfey et al., 2003; Van’t Wout et al., 2006; Koenigs & Tranel, 2007). Punishing the proposers for behaving unfairly was found to be rewarding for the responders, because it compensates for the negative affective reaction to being treated unfairly (De Quervain et al., 2004). When responders are motivated to maximize their gain, they tend to adopt a more deliberative reasoning allocating more cognitive resources to the decision process (e.g., Fabre et al., 2015; Halali et al., 2014). This increased cognitive effort is, at least partly, dedicated to the reappraisal − i.e., a cognitive change involving the reinterpretation of an event’s meaning to alter its emotional impact (Gross, 1998) − of the negative affective response to unfair shares and is associated with less predictable decisions (e.g., Grecucci et al., 2013; Van’t Wout et al., 2010). Moreover, depleting responders’ cognitive resources was found to affect the reappraisal of the negative affective response to unfair offers and trigger higher rejection rates of unfair shares, (Halali et al., 2014).

### 1.3. The ultimatum game and the framing effect

Framing manipulations are expected to modulate the way the offers are perceived by the players and affect their behavior accordingly (Kahneman & Frederick, 2007). In previous studies, the framing manipulation consisted in manipulating the way the proposers’ behavior is presented to the participants playing as responder, with the offers framed either positively with the focus being put on the money received by the responder or negatively for the responders’ point of view, focusing on the money removed from the responder (Sarlo et al., 2013; Tomasino et al., 2013). Two previous studies investigated the impact of the offer frame manipulation on responders (Sarlo et al., 2013; Tomasino et al., 2013). In the study of Sarlo and colleagues (2013) both skin conductance and heart rate were recorded. The results showed an effect of the offer frame manipulation at the physiological level in both male and female participants and at the behavioral level in male participants only. Specifically, higher rejection rates of mid-value offers (i.e., 3€ out of 10€) in male participants; and an increase in both heart rate and skin conductance in overall participants were observed in the negative frame compared to the positive frame, suggesting a major defense response (i.e., affective reaction) to negatively framed offers. The *f*MRI study of Tomasino and colleagues (2013) aimed at investigating the neural mechanisms underpinning the offer framing effect, using a paradigm similar to the one used in the study of Sarlo and colleagues (2013). Although the authors lacked to find a main effect or a significant interaction involving frame at the behavioral level, neuroimaging data revealed an increase in neural activity in the right rolandic operculum/insular cortex and the anterior cingulate cortex (ACC) for accepting versus rejecting offers in the negative frame, as compared to the positive frame. The activation of the operculo-insular cortex was interpreted to signal this deviation from participants’ own expected behavior (i.e., violation of social norms). The authors suggested that the ACC activation may reveal the major conflict between cognitive and emotional motivations triggered by accepting the offers when the proposer takes the money (i.e., negative frame) than when s/he offers it (i.e., positive frame). Taken together, the results of these two studies are far from straightforward, making it difficult to status on the impact of the way the offers’ framing impacts the responders’ behavior.

### 1.4. The Present Study

The present study aimed at investigating whether offer frame manipulations modulated the acceptance behavior of responders playing the ultimatum game. Experiment 1 was an ERP study reproducing the studies of Tomasino and colleagues (2013) and Sarlo and colleagues (2013) where participants played the repeated one-shot ultimatum game as responders (i.e., each new trial played with a different proposer). The aim of this experiment was twofold: 1) determining whether or not the way the offers are framed in the ultimatum game modulates the responders’ decision-making; and 2) investigating the Event-Related Potentials (ERPs) associated with the offers depending on their frame to understand how offer frame manipulations impact responders’ decision process. Experiments 2 and 3 were on-line experiments. The aim of Experiment 2 was to determine whether offer frame manipulations affect responders diversely when the ultimatum game is played only once and not repeatedly as in Experiment 1. In Experiment 2, the framing manipulation was the same as Experiment 1, but participants had to decide to accept or reject only one economic offer from a unique proposer (i.e., one-shot ultimatum game). In Experiment 3, we tested the assumption that increasing the responders’ cognitive load (i.e., making the ultimatum game statement more complex) would trigger a framing effect in responders. Participants played a one-shot ultimatum game where two frame manipulations were applied (an offer frame manipulation and an action frame manipulation) increasing the complexity of the game rules.

## 2. EXPERIMENT 1

Event Related Potentials (ERPs) allow the investigation of attentional, cognitive and emotional processes underpinning economic decision-making with a high temporal resolution (e.g., Fabre et al., 2015). In recent years, this technique has been applied to the study of the processes underpinning framing effects in consumer studies (Jin et al., 2017; Ma et al., 2018), life and death decision studies (Ma et al., 2015; Ma et al., 2012) or gambling studies (Ma et al., 2015; Yu & Zhang, 2014). However, to our knowledge no ERP study was conducted to investigate the responders’ electrophysiological brain responses to the way the offers were framed in the ultimatum game. Experiment 1 aimed at filling this gap in the literature.

In the previous ERP studies investigating the impact of framing effect, the information of interest was framed and presented using either numbers (Ma et al., 2012, 2015), charts (Jin et al., 2017), or a mix of pictures and amounts of money (Ma et al., 2018); or both words and pie charts (Yu & Zhang, 2014). Four of these studies found greater N2/Feedback Related Negativity (FRN) amplitudes when the information of interest was framed negatively compared to when it was framed positively (Jin, et al., 2017; Ma et al., 2012, 2015; Yu & Zhang, 2014). The FRN is a negative component observed at frontal-central sites that reaches its maximum in the 200-300 ms time windows after the stimulus onset (for an overview, see Walsh & Anderson, 2011). This component was found to be sensitive to the valence of the outcomes, with greater FRN responses to unfavorable outcomes than to favorable ones (e.g., Fabre et al., 2015; Gehring & Willoughby, 2002; Hajcak et al., 2006; Polezzi et al., 2008). Consequently, similar outcomes appear to have been classified as less favorable when presented in the negative frame than in the positive frame. Two studies also found greater Late Positive Potential (LPP) amplitudes in response to positively framed items than to negatively-framed items (Jin et al., 2017; Ma, et al., 2018). The LPP is a positive component maximal at central-parietal sites and peaking at in the 400 - 800 ms time window after the stimulus onset (Schupp et al., 2000). The LPP component was found to be sensitive to motivationally relevant stimuli and to reflect the deliberative processing of the stimulus significance (Olofsson et al., 2008), suggesting that participants may have been more motivated to purchase positively-framed than negatively-framed items (Jin et al., 2017; Ma et al., 2018).

While the analysis of the brain electrophysiological correlates (i.e., ERPs) was found to be an effective tool to understand the processes underpinning framing manipulations, no ERP study was conducted to investigate the brain responses to offer frame manipulations in the ultimatum game. The aim of Experiment 1 was twofold: 1) replicating previous ultimatum game studies to determine whether or not the way the offers are framed in the ultimatum game modulates the responders’ decision-making and 2) investigating the ERPs observed in response to framed-offers, to improve our understanding of the processes underpinning the responders’ decision-making in response to offer frame manipulations. Participants played a repeated one-shot ultimatum game (i.e., a different proposer for each offer) as responders. The offers were either framed positively (the proposer says “I give you 1€/3€/5€”) or negatively (the proposer says “I keep for myself 9€/7€/5€”).

In line with previous studies, we expected the responders to accept more frequently fair offers (i.e., 5€ out of 10€) than mid-value offers (i.e., 3€ out of 10€) than unfair offers (i.e., 1€ out of 10€; Camerer, 2003). Two main ERPs are usually observed in response to the offers in the ultimatum game: the FRN component and the P300 component (e.g., Fabre et al., 2015). Greater FRN amplitudes are usually found in response to non-fair offers (i.e., both mid-value and unfair offers) than to fair offers (Alexopoulos et al., 2012, 2013; Fabre et al., 2015; Polezzi et al., 2008; Qu et al., 2013; Wu et al., 2011, 2012), and interpreted as reflecting the responders’ internal conflict between their desire to reject non-fair offers to punish the proposers for violating social norms (i.e., non-respect of the equity) and accepting every offer to maximize their final outcome (e.g., Fabre et al., 2015). The P300 is a positive component occurring in the 300–600 ms time window after the stimulus onset and maximal at central-parietal sites (Luck & Kappenman, 2011). This component was found to underpin the allocation of attentional resources to the decision-making process (e.g. Gray et al., 2004; Falco et al., 2019) and to reflect the decision maker’s motivational state (Yeung & Sanfey, 2004; Nieuwenhuis et al., 2005). In line with literature, we predicted to observe 1) greater FRN amplitudes in response to non-fair offers compared to fair offers; and 2) greater P300 amplitudes in response to fair offers than to unfair offers, reflecting responders’ greater motivation to accept fair offers than unfair offers (in line with Qu et al., 2013; Wu et al., 2012).

Due to contradictory results of previous ultimatum game studies (Sarlo et al., 2013; Tomasino, et al., 2013), we were not able to predict whether the offer frame manipulation would modulate participants’ decision behavior. At the difference of the previous ERP studies investigating framing effect, in the present experiment the information of interest (i.e., the offers) was presented in sentence form. Hence, we made the hypothesis that offer frame manipulations would either impact the process of the offer’s value assessment at an early stage (i.e., FRN and P300); and/or the process of its integration in the sentence context at a later stage (i.e., N400 and P600). In the case the frame manipulation affects the processing of the offers’ value, modulations of the FRN and/or the P300 components might be observed. We predicted to observe greater FRN amplitudes in response to the offers in the negative frame than in the positive frame, since focusing on the money removed from them may be more contentious for the participants (Ma et al., 2012; Ma et al., 2015). We also predicted to observe greater P300 amplitudes in response to the offers in the positive frame than in the negative frame (especially to unfair ones), reflecting the greater motivation of participants toward positively framed offers than to negatively framed offers. In the case the frame manipulation affects the integration of the offers in the context of the sentence, we predicted that we might observe modulations of two components known to underpin language processing: the N400 component and the P600 component. The N400 is a negative going component peaking around 400 ms in the 200-600 ms time window and its maximal is found over central-parietal sites with a slight right hemisphere bias (for a review, see Kutas & Federmeier, 2011). First observed by Kutas and Hillyard (1980), the N400 component is sensitive to the contextual relationship between an eliciting word and its preceding, and was found in response to both conceptual and emotional incongruities (Kutas & Federmeier, 2011; Chen et al., 2013). In absence of semantic anomalies, some study reported greater N400 responses to emotional words than to neutral words (Bayer, et al., 2010; Holt, et al., 2009; Ding et al., 2015); and to negative words than to positive words (De Pascalis et al., 2009; Wang et al., 2013). The P600 (sometimes called Late Positivity Component) is a positive going component peaking approximately 600 ms after stimulus onset and maximal at central-parietal sites (Bornkessel-Schlesewsky & Schlesewsky, 2008). This component is thought to be part of the P300 family and to underpin the attention-related stimulus reevaluation and the working memory update (Sassenhagen et al., 2014). Greater P600 amplitudes were found to reflect both syntactic (Kaan et al., 2000) and semantic (Van Herten et al., 2005) reprocessing of the information within the context (Dorjee et al., 2015). In the case frame manipulations affect the integration of the offers in the sentence context, we predicted to observe greater N400 amplitudes in response to the offers in the negative frame than in the positive frame, reflecting the greater negative affective response to the offers in the negative frame than in the positive frame. If responders are motivated to maximize their outcome, they may engage in a reevaluation of negatively framed offers (especially unfair ones) to appraise the negative emotional reaction associated with these offers. In this case, we might observe greater P600 responses to the offers in the negative frame than in the positive frame, this effect being even more pronounced for unfair offers.

### 2.1. Material and Methods

#### 2.1.1. Participants

34 Italian students at Modena University, Polytechnic of Turin or Polytechnic of Milan (9 females, *M* = 25 years old **±** 4, age range from 19 to 30 years old) participated in this study. All were right-handed and had normal or corrected-to-normal vision. None of the participants reported a history of prior neurological disorder.

#### 2.1.2. Ethics statement

All participants were informed of their rights, that they could withdraw from the experiment at any time and gave written informed consent for participation in the study, according to the Declaration of Helsinki. The research was carried out fulfilling ethical requirements in accordance with the standard procedures of the University of Modena and Reggio Emilia.

#### 2.1.3. Procedure

We used an experimental procedure similar to those employed in previous studies that investigated the framing effect using the ultimatum game paradigm (Sarlo et al., 2013; Tomasino, et al., 2013). Participants played a repeated one-shot ultimatum game as responders. The instructions were presented in a written form before the experiment started. The instructions can be subsumed as follows. Before conducting the game, participants were told that they were playing as responder against 230 real proposers that would offer to split 10 €. The participant’s task consisted in accepting or rejecting the offers that were proposed to them. In the case they accepted the offer; the money would be split as proposed by the proposer. However, if they rejected the offer, both players would lose their money. They were told that previous participants played as proposers and were asked to indicate how much money they would offer a responder and how they wanted to present the offer (i.e., positive versus negative frame). These offers were filled into the experiment’s database and used in the game. Participants were informed that they would receive a payment consisting of the 1% of the money they accepted in the game at the end of the experiment. Only at the end of the experiment, the participants were proposed to swap the money they won for a course credit.

Participants were seated comfortably in a darkened sound-attenuated room. Stimuli were presented in white uppercase letters (Courier font, size 14) against a black background on a high-resolution computer that was positioned at eye level 70 cm in front of the participant. A fixation point (+) appeared in the middle of the screen and stayed there until participants pressed the space bar to start a trial. Then a blank screen was displayed for 300 ms. Then *Participant x says* (“Partecipante x dice”; with x ranging from 1 to 230) followed *by I give you* (“Ti do”) or *I keep for myself* (“Mi tengo”) appeared on the screen for 700 ms. Each presentation was followed by a blank screen for 300 ms. Finally, the offer appeared and remained on the screen until the participants made their decision. Half of the participants pressed the right button to accept the offer (i.e., corresponding to the M on the keyboard) and the left button to reject the offer (i.e., corresponding to the C on the keyboard). The other half did the reverse. Each response was followed by a 1000 ms blank screen.

Before starting the experiment, participants took part in a short training session consisting of 10 trials. Each experimental session consisted in 230 randomized trials, including 210 experimental trials [3 (Offer: 1€, 3€, 5€) × 2 (Frame: negative frame “I keep for myself”, positive frame “I give you”) × 35 repetitions], yielding a total of 105 offers for each frame condition and 20 trials of no interest (5 offers with 2€ and 5 offers with 4€ proposed to the responder), were included for each frame condition in order to represent the full range of offers that the proposers may make (Sarlo et al., 2012; Tomasino et al., 2013).

#### 2.1.4. Electroencephalograph (EEG) recordings

EEG was amplified and recorded with a BioSemi ActiveTwo system from 30 Ag/AgCl active electrodes (http://www.biosemi.com) mounted on a cap and placed on the scalp according to the International 10–20 System (FP1, FP2, AF3, AF4, F7, F3, Fz, F4, F8, FC5, FC1, FC2, FC6, CP5, CP1, Cz, CP2, CP6, P7, P3, Pz, P4, P8, T7, T8, PO3, PO4, O1, Oz, O2) plus two sites below the eyes for eye movement monitoring. Two additional electrodes placed close to Cz, the Common Mode Sense [CMS] active electrode and the Driven Right Leg [DRL] passive electrode, were used to form the feedback loop that drives the average potential of the participant as close as possible to the AD-box reference potential (Van Rijn, et al., 1990). Skin-electrode contact, obtained using electro-conductive gel, was monitored, keeping voltage offset from the CMS below 25 mV for each measurement site. All the signals were DC-amplified and digitalized continuously with a sampling rate of 512 Hz with an anti-aliasing filter with 3 dB point at 104 Hz (fifth order sinc filter); no high-pass filtering was applied online. The triggering signals to each word onset were recorded on additional digital channels. EEG data were analyzed using EEGLAB v.13.6.5b open-source software (Delorme & Makeig, 2004) and ERPLAB toolbox (Lopez-Calderon, & Luck, 2014) on Matlab 2016b. EEG data were off-line re-referenced to the average activity of the two mastoids and band-pass filtered (0.1 – 40 Hz, 12 dB/octave), given that for some subjects the low-pass filter was not effective in completely removing the 50 Hz artifact. An independent component analysis was performed to isolate eye-blinks and movements related artifacts that were removed from the signal. Then, 1000 ms epochs containing the ERP elicited by the offers were extracted, starting with 200 ms prior to the onset of the offer. Segments with excessive blinks or including artefacts (such as excessive muscle activity) were eliminated off-line before data averaging. One participant was excluded from the analyses due to the high number of rejected epochs (> 40 %). The lost data (due to artefacts) of the remaining 33 participants were about 9 %. A 200 ms pre-stimulus baseline was used in all analyses.

### 2.2. Results

#### 2.2.1. Behavioral results

Two 2 (Offer Frame: negative frame; positive frame) x 3 (Offer: 1€; 3€; 5€) repeated analyses of variance (ANOVAs) were conducted on both acceptance rates and response times. LSD post-hoc tests were carried out to further examine significant effects (α < .05).

##### Acceptance rates

The analysis showed a significant main effect of offer [*F* (2, 64) = 136.49, *p* < .001, η*p2* = .81; see Figure 1.A.]. Participants accepted significantly more frequently fair offers (*M* = 96.88 %*, SD* = 12.95) than both mid-value (*M* = 31.21%*, SD* = 40.33, *p* < .001) and unfair (*M* = 6.83 %*, SD* = 20.99, *p* < .001) offers; and mid-value offers than unfair offers (*p* < .001). The main effect of offer frame [*F* (1, 32) = .01, *p* = .94, η*p2* = .00] and the Offer Frame x Offer interaction [*F* (2, 64) = 2.33, *p* = .11, η*p2* = .07] did not reach significance.

**Figure 1.**
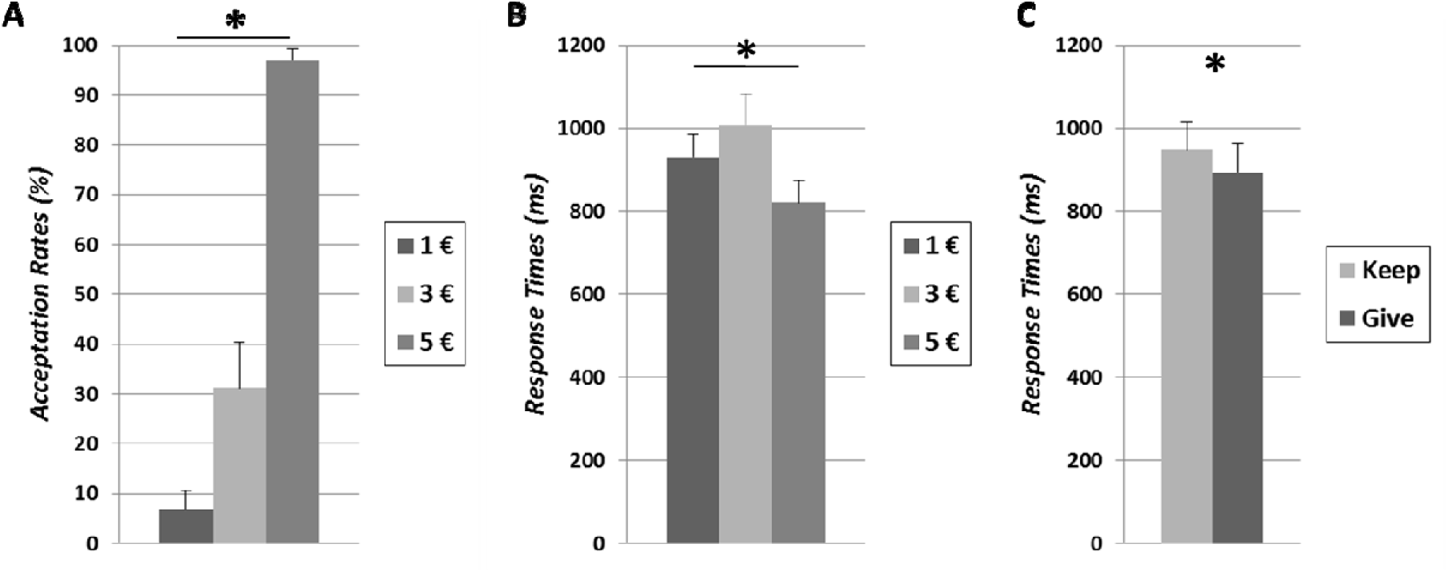
***A)*** Acceptances rates and ***B)*** response times for 1€ (dark grey), 3€ (light grey) and 5€ offers (mid-grey). ***C)*** Response times for offers in the positive frame (dark grey) and in the negative frame (light grey). Error bars represent standard errors.

##### Response times

The analysis revealed a significant main effect of offer [*F* (2, 64) = 17.50, *p* < .001, η*p2* = .35; see Figure 1.B.], in that participants were faster in answering fair offers (*M* = 821 ms, *SD* = 226) than both mid-value (*M* = 1009 ms, *SD* = 310; *p* < .001) and unfair offers (*M* = 931 ms, *SD* = 238; *p* < .01.); and unfair offers than mid-value offers (*p* < .05). The analysis also revealed a main effect of offer frame [*F* (1, 32) = 12.71; *p* < .001; η*p2* = .28; see Figure 1.C] with longer response times to the offers in the negative frame than in the positive frame (respectively, *M* = 948 ms, *SD* = 272; *M* = 893 ms, *SD* = 267). The Offer Frame x Offer interaction did not reach significance [*F* (2, 64) = .35, *p* = .70, η*p2* = .01].

#### 2.2.2. Brain electrophysiological results

Overall participants were presented with 35 trials per condition (210 trials in total). After removing the artefacts, each participant had on average 31.9 trials (*SD* = 2.8) per condition for EEG averaging. The FRN and the P300 components were assessed in terms of mean amplitude at Fz, Cz and Pz electrodes respectively in the 220 – 300 ms and 300 – 380 ms time windows (Fabre et al., 2015). Due to the important amplitude differences observed before the occurrence of the N400 and the P600 components, these two components were assessed in terms of peak-to-peak amplitudes (Falco et al., 2019). For the N400 component, peak-to-peak amplitudes were calculated by subtracting the peak amplitudes measured in the 300 – 380 ms (positive peak) and 370 – 430 ms (negative peak) time windows. For the P600 component, peak-to-peak amplitudes were calculated by subtracting the peak amplitudes measured in the 370 – 430 ms (negative peak) and the 400 – 650 ms (positive peak) time windows. For each component, a 2 x 3 [Offer Frame (negative frame; positive frame) x Offer (1€, 3€, 5€) x Electrode (Fz, Cz, Pz)] repeated measures ANOVA was conducted on the amplitude measurements. LSD post-hoc tests were carried out to further examine significant effects (α < .05).

##### FRN component

The ANOVA analysis revealed a main effect of electrode [*F* (2, 64) = 30.50, *p* < .001, η*p2* = .49] with greater FRN responses at Fz electrode (*M* = 3.15 µV, *SD* = 4.14, *p_s_* < .001) than at Cz (*M* = 5.62 µV, *SD* = 3.93) and Pz (*M* = 5.01 µV, *SD* = 5.01) electrodes. The FRN responses were not significantly different at Cz and Pz electrodes (*p* = .17). The main effects of offer frame [*F* (1, 32) = 3.87, *p* = .058, η*p2* = .11] and offer [*F* (2, 64) = 2.33, *p* = .11, η*p2* = .07]; and the Offer Frame x Offer [*F* (2, 64) = 1.29, *p* = .28, η*p2* = .04], the Offer Frame x Electrode [*F* (2, 64) = 1.30, *p* = .28, η*p2* = .04], the Offer x Electrode [*F* (4, 128) = 1.93, *p* = .11, η*p2* = .06] and the Offer Frame x Offer x Electrode [*F* (4, 128) = 1.16, *p* = .33, η*p2* = .04] interactions did not reach significance.

##### P300 component

The ANOVA analysis revealed a main effect of electrode [*F* (2, 64) = 9.97, *p* < .001, η*p2* = .24] with lower amplitudes measured at Fz (*M* = 4.43 µV, *SD* = 3.52) than at both Cz (*M* = 5.47 µV, *SD* = 3.70, *p* < .001) and Pz (*M* = 5.86 µV, *SD* = 4.10, *p* < .001) electrodes. The amplitudes at Cz and Pz were not significantly different (*p* = .24). The analysis also revealed a main effect of offer [*F* (2, 64) = 5.32, *p* < .01, η*p2* = .14; ; see Figure 2.A.] with greater amplitudes in response to fair offers (*M* = 5.91 µV, *SD* = 4.01) than to both unfair (*M* = 4.83 µV, *SD* = 3.59, *p* < .05) and mid-value offers (*M* = 5.03 µV, *SD* = 3.79, *p* < .01). The main effects of offer frame [*F* (1, 32) = 1.70, *p* = .20, η*p2* = .05]; and the Offer Frame x Offer [*F* (2, 64) = 1.39, *p* = .26, η*p2* = .04], the Offer Frame x Electrode [*F* (2, 64) = .39, *p* = .68, η*p2* = .01], the Offer x Electrode [*F* (4, 128) = 1.35, *p* = .26, η*p2* = .04] and the Offer Frame x Offer x Electrode [*F* (4, 128) = .45, *p* = .77, η*p2* = .01] interactions did not reach significance.

**Figure 2.**
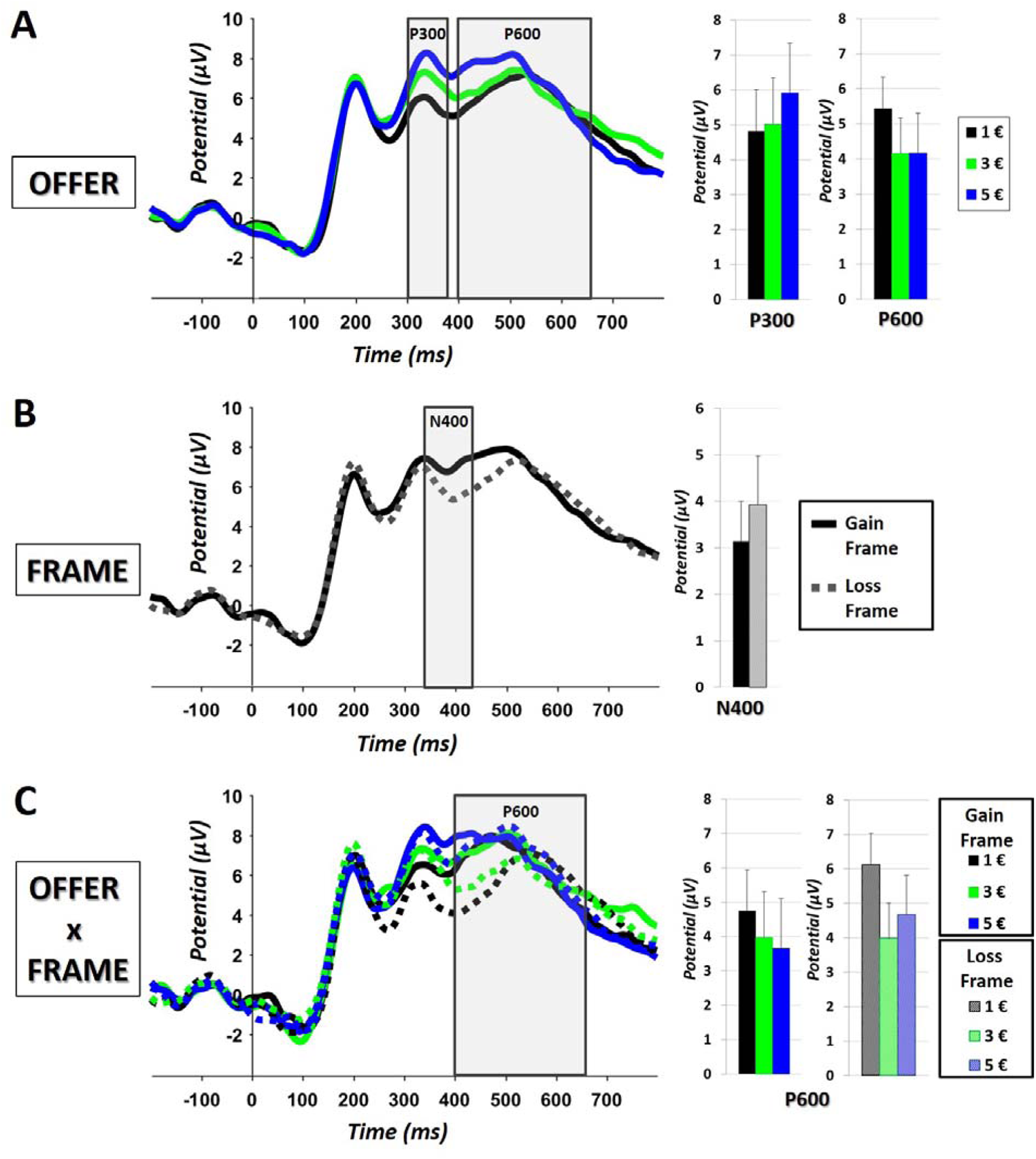
Grand average event-related brain potential waveforms observed at Cz electrode elicited by ***A)*** unfair offers (black line), mid-value offers (green line) and fair offers (blue line); ***B)*** offers in the positive frame (solid lines) in the negative frame (dashed lines); ***C)*** unfair offers (black line), mid-value offers (green line) and fair offers (blue line) in the positive frame (solid lines) and in the negative frame (dashed lines). Negative is plotted down. Zeros on the timeline indicate the onset of the offer.

##### N400 component

The analysis also revealed a main effect of offer frame [*F* (1, 32) = 13.87, *p <* .001, η*p2* = .30; see Figure 2.B.] with a greater N400 amplitudes in response to the offers in the negative frame (*M* = 3.93 µV, *SD* = 1.05) than in the positive frame (*M* = 3.14 µV, *SD* = .87). The main effects of electrode [*F* (2, 64) = .99, *p* = .38, η*p2* = .03] and offer [*F* (2, 64) = .13, *p* = .88, η*p2* = .00]; and the Offer Frame x Offer [*F* (2, 64) = .35, *p* = .70, η*p2* = .01], the Offer Frame x Electrode [*F* (2, 64) = .33, *p* = .72, η*p2* = .01], the Offer x Electrode [*F* (4, 128) = 1.58, *p* = .18, η*p2* = .05] and the Offer Frame x Offer x Electrode [*F* (4, 128) = .89, *p* = .47, η*p2* = .03] interactions did not reach significance.

##### P600 component

The analysis revealed a main effect of electrode [*F* (2, 64) = 3.81, *p <* .05, η*p2* = .11] with greater amplitudes measured at Cz (*M* = 5.08 µV, *SD* = 3.41) than at both Fz (*M* = 4.24 µV, *SD* = 3.07, *p* < .05) and Pz (*M* = 4.43 µV, *SD* = 3.47, *p* < .05) electrodes. No difference in amplitude was found between Fz and Cz (*p* = .56). The analysis also revealed a main effect of offer [*F* (2, 64) = 5.70, *p <* .01, η*p2* = .15; see Figure 2 A], with greater amplitudes in response to unfair offers (*M* = 5.43 µV, *SD* = 3.17) than to both mid-value (*M* = 4.15 µV, *SD* = 3.20, *p* < .01) and fair (*M* = 4.17 µV, *SD* = 3.48, *p* < .01 offers. No difference in amplitude was found between fair and mid-value offers (*p* = .99). The analysis also revealed a significant Offer Frame x Offer interaction [*F* (2, 64) = 3.15, *p <* .05, η*p2* = .09; see Figure 2. C.], with greater amplitudes in response to unfair offers (*M* = 6.12 µV, *SD* = 3.23) than to mid-value offers (*M* = 3.98 µV, *SD* = 3.07, *p* < .001) and fair offers (*M* = 4.67 µV, *SD* = 3.75, *p* < .01) in the negative frame. Greater amplitudes were also found in response to unfair offers (*M* = 4.75 µV, *SD* = 2.96) than fair offers (*M* = 3.67 µV, *SD* = 3.14, *p* < .05) in the positive frame; and to unfair offers in the negative frame than to unfair offers (*p* < .01), mid-value offers (*M* = 4.32 µV, *SD* = 3.32, *p* < .001) and fair offers (*p* < .001) in the positive frame. The main effect of offer frame [*F* (1, 32) = 3.05, *p* = .09, η*p2* = .09]; and the Offer Frame x Electrode [*F* (2, 64) = .06, *p* = .94, η*p2* = .00], the Offer x Electrode [*F* (4, 128) = .41, *p* = .80, η*p2* = .01] and the Offer Frame x Offer x Electrode Offer [*F* (4, 128) = .28, *p* = .89, η*p2* = .01] interactions did not reach significance.

### 2.3. Discussion

The aim of Experiment 1 was twofold. First, to investigate whether the way an offer is framed modulates the decision-making of responders playing the ultimatum game; and second, to ascertain the ERPs associated with the offer frame manipulations in the ultimatum game. Participants played a repeated one-shot ultimatum game as responders where the offers were either framed in a negative or a positive way.

Consistent with the literature, we found that the responders were more likely to accept fair offers than mid-value and unfair offers; and mid-value offers than unfair offers (Camerer, 2003). Responders also took longer to answer non-fair offers (i.e., both mid-value and unfair offers) than fair offers. Greater P300 amplitudes were found in response to fair offers than to both mid-value and unfair offers, reflecting the responders’ greater motivational significance to fair shares than to non-fair shares (in line with Wu et al., 2012; Qu et al., 2013). This result suggests that at this point of the process, the economic value of the offers was retrieved and indexed in both frames.

In line with the results of the study of Tomasino and colleagues (2013), no effect of the offer frame manipulation was found on the responders’ decision behavior in the present experiment. However, participants were longer to answer the offers when they were framed negatively than positively, suggesting that it may have been more complex to deal with the offers in the negative frame than in the positive frame (in line with Sarlo et al., 2013). Some interesting results were found at the electrophysiological level. The frame manipulation did not impact the components underpinning the evaluation of the outcome (i.e., FRN and P300 components) but modulated the amplitude of two components reflecting language processing: the N400 component and the P600 component. Greater N400 amplitudes were found in response to the offers in the negative than in the positive frame. Previous studies found greater N400 amplitudes in response to stimuli triggering negative emotional responses (e.g., Bayer et al., 2010), suggesting that negatively framed offers triggered a greater negative affective response than positively framed offers. Finally, greater P600 amplitudes were found in response to 1) unfair offers than to both mid-value and fair offers; and 2) unfair offers in the negative frame than in the positive frame. The P600 component is interpreted as reflecting the reevaluation/reprocessing of the information within the context (Dorjee et al., 2015). Consequently, the present results suggest that participants intensively reprocessed unfair offers, especially in the negative frame, presumably aiming at reappraising the negative affective response to these offers. This reprocessing appears to have been successful since the way the offers were framed did not impact participants’ decision behavior (Miu & Crişan, 2011).

Taken together, the results of the present study support the cognitive–affective tradeoff model (Gonzalez et al., 2005). Negatively framed offers triggered a negative affective response in responders (i.e., N400) compared to positively framed offers. The reappraisal of this negative affective reaction was associated with a costly cognitive reprocess of the information reflected by longer response times and greater P600 amplitudes (in line with Miu & Crişan, 2011). Participants of both the present study and the study of Tomasino and her colleagues (2013) appear to have retrieved enough cognitive resources to reappraise the negative affective response to negatively framed offers, explaining why a framing effect was not always observed in these ultimatum game studies. The framing effect may arise when the affective response is too strong to be reappraised and/or insufficient cognitive resources are retrieved to reappraise the affective response in the negative frame.

Therefore, there may be two main possible explanations to the fact that no framing effect was found in the present experiment and the study of Tomasino and colleagues (2013). On the one hand, it is possible that negatively framed offers did not trigger a negative affective response strong enough to affect the responders’ behavior, either because the ultimatum game is associated with a low emotional resonance compared to life-and-death decision problems for instance (for an overview see Costa et al., 2014) and/or because repeated measurements triggered an habituation effect and lessened the affective response to negatively framed offers throughout the experiment (e.g., Emmelkamp, 2004). On the other hand, the ultimatum game being a quite simple task with certain outcomes for the responders, participants may have easily retrieved the cognitive resources necessary to reappraise the negative affective response to the offers in the negative frame.

Experiments 2 and 3 were designed to test these two hypotheses. Experiment 2 aimed at investigating the impact that repeated measures may have had on the results of Experiment 1. Experiment 3 aimed at testing whether the absence of framing effect could be attributed (at least partly) to the relative simplicity of the ultimatum game.

## 3. EXPERIMENT 2

Repeated exposure to aversive stimuli can decrease both affective and physiological responses to these stimuli (i.e., habituation; Emmelkamp, 2004). In both Experiment 1 of the present study and the study of Tomasino and colleagues (2013), participants were exposed repeatedly to the offers (i.e., 35 times to each type of offers in Experiment 1). The repeated exposure to the negatively framed offers could have triggered a habituation in participants and lessened the negative affective response to these offers throughout the experiment, resulting in an absence of framing effect. Experiment 2 aimed at testing this assumption. In this experiment, the offer frame manipulation was identical to Experiment 1, but participants played a one-shot ultimatum game as responders, where they had to decide whether to accept or reject only one offer from one unique proposer. In line with the literature, we predicted that responders would accept more frequently fair offers (i.e., 5€ out of 10€) than mid-value offers (i.e., 3€ out of 10€) than unfair offers (i.e., 1€ out of 10€; Camerer, 2003). In case the repeated exposure to negatively framed offers had triggered a significant habituation effect in Experiment 1, we expected to observe an offer framing effect with participants accepting more frequently positively framed offers than negatively framed offers.

### 3.1. Material and Methods

#### 3.1.1. Participants

769 Italian participants (503 females; mean age: 32 years ± 11) took part in Experiment 2. They received no compensation for their participation in the study.

#### 3.1.2. Ethics statement

All participants were informed of their rights, that they could withdraw from the experiment at any time and gave written informed consent for participation in the study, according to the Declaration of Helsinki. The research was carried out fulfilling ethical requirements in accordance with the standard procedures of the University of Modena and Reggio Emilia.

#### 3.1.3. Procedure

When clicking on the website link, participants were directed to the front page of the questionnaire made on Qualtrics ©. They were explained that the experiment would take less than three minutes and were asked to click on the icon “participate” to continue to the experiment. After clicking on the “participate” icon they were directed to the instruction page. They were asked to attentively read the instructions before starting the experiment. They were explained that the anonymity of their personal data was respected as no IP address would be collected during the experiment. They were also asked to participate in the experiment only once. Participants were informed that they were free to leave the experiment at any time (i.e., incomplete responses would not be accounted for). They were also explained that once they had finalized the experiment by clicking on the “finalize” icon, it would no longer be possible to modify their response. They were informed that they could click on the “contact” icon to contact the researchers in charge of the experiment and communicate any information they wanted. By clicking a second time on the icon “participate” they acknowledged that they had read and understood the instructions and accepted the terms of the experiment. In the third website page, the ultimatum game problem was presented as follows: “*You are about to play a game called the ultimatum game. In this game, a proposer is given a fix amount of money (i.e., 10*€*) and has to choose how to divide it with a second player called responder. The responder can either accept the division, in which case the amount of money gets divided as proposed by the proposer, or refuse it, in which case neither of the two gets any money*”. For the 386 participants (250 females) who were tested in the positive frame, the final sentence of the problem was: “*Playing as responder, what would you do if the proposer gave you X*€ *on 10*€*?”* (with X = 1€, 3€ or 5€)*”*. For the 383 participants (254 females) who were tested in the negative frame, the final sentence was: “*Playing as responder, what would you do if the proposer kept Y*€ *on 10*€*?”* (with Y = 9€; 7€ or 5€). The participants’ task was to click either on the icon “accept” or on the icon “reject”. Afterwards, they were sent to a fourth website page and asked to enter their age and biological gender and then click on the icon “Terminate” to validate their choice and complete the experiment. Finally, they were then thanked for participating in the study.

### 3.2. Results

A 2 x 2 x 3 (Offer Frame [positive frame, negative frame] x Participants’ Gender [male, female] x Offer [1€; 3€; 5€]) binary logistic regression was performed on the decision to either accept or reject the offer. A manual stepwise analysis was performed to remove non-significant interactions from the model. The analysis revealed a main effect of offer [*B* (SE) = - 1.809 (.221), CI _(95%)_ = (- 2.242, - 1.378), Wald χ*²* (2) = 67.819, *p* < .001; see Figure 3] with 5€ offers (*M* = 86.19 %, *p_s_ <* .001) predicting for a higher acceptance rate than 1€ (*M* = 50.78 %) and 3€ (*M* = 61.20 %) offers; and 3€ offers predicting for a higher acceptance rate than 1€ (*p <* .05). The main effect of offer frame [*B* (SE) = .103 (.226), CI _(95%)_ = (- .212, .418), Wald χ*²* (1) = .414, *p* = .52] and participants’ gender [*B* (SE) = .128 (.168), CI _(95%)_ = (- .201, .457), Wald χ*²* (1) = .585, *p* = .45] did not reach significance.

**Figure 3.**
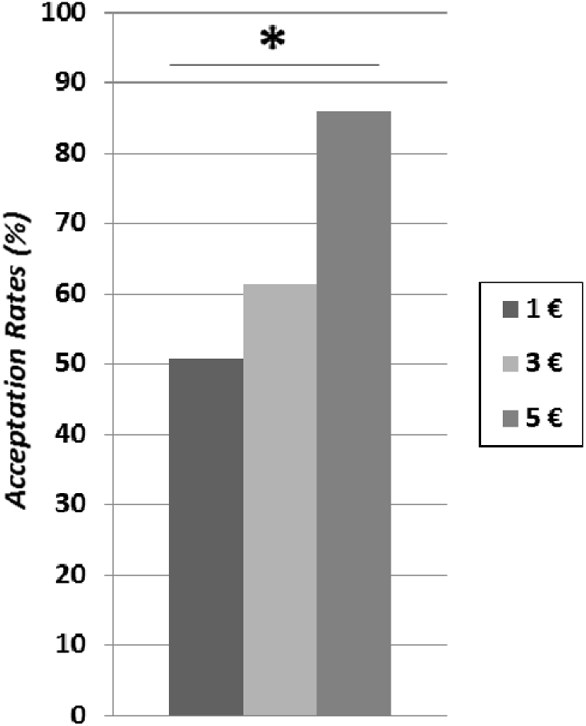
Acceptances rates for 1€ (dark grey), 3€ (mid-grey) and 5€ offers (light grey).

### 3.3. Discussion

Experiment 2 aimed at investigating whether the absence of framing effect in Experiment 1 of the present study and the study of Tomasino and colleagues (2013) could be attributed to their repeated-measure design. Participants played a one-shot ultimatum game as responders and were proposed one unique offer. In line with the literature, the results showed a significant effect of the offers’ value, with fair offers predicting for a higher acceptance rate than both mid-value and unfair offers; and mid-value offers predicting for a higher acceptance rate than unfair offers. Interestingly, acceptance rates of both unfair and mid-value offers were much higher in the present experiment where participants played a one-shot ultimatum game (unfair offers: 50.78 %; mid-value offers: 61.20 %) than in Experiment 1 where participants played a repeated one-shot ultimatum game (unfair offers: 6.83 %; mid-value offers: 31.21%). Various studies showed that when responders played a repeated ultimatum game as in Experiment 1 of the present research, their behavior adapted along trials (Avrahami et al., 2013; Cooper & Dutcher, 2011). They were found to demand lower shares at the beginning of the experiment, but their behavior quickly became more egalitarian (i.e., they demanded shares close to 50% of the amount of money). The results of these studies are in line with the results of the present study and explain why responders were more likely to accept non-fair offers when playing a one-shot ultimatum game (Experiment 2) than a repeated ultimatum game (Experiment 1). In line with Experiment 1, the results of the present experiment showed no effect of the offer frame manipulation on responders’ decision behavior. These results suggest that it is unlikely that the absence of offer framing effect in Experiment 1 was due to a decrease in negative affective response to negatively-framed offers over time (i.e., a habituation effect) triggered by a repeated exposure to this type of offers.

## 4. EXPERIMENT 3

Cognitive processes play an important role in economic decision-making (e.g., Westbrook & Braver, 2015). In the ultimatum game, depleting responders’ cognitive resources was found to increase the rejection rate of unfair shares, seemingly because the lack of cognitive resources did not allow the reappraisal of the negative affective response to being treated unfairly by the proposer (Halali et al., 2014). While both physiological (Sarlo et al., 2013) and neural (Experiment 1 of the present study; Tomasino et al., 2013) correlates of offer frame manipulations have shown that a greater negative affective response was found in response to the offers in the negative than in the positive frame, an offer framing effect (at a behavioral level) was found in neither the study of Tomasino and colleagues (2013) nor Experiments 1 and 2 of the present study. The brain electrophysiological results of Experiment 1 suggest that the reappraisal of the negative affective response to the offers in the negative frame underpinned by a costly cognitive process prevented the emergence of an offer framing effect (in line with Kahneman & Frederick, 2007; Miu & Crisan, 2011).

Many studies have investigated the involvement of cognitive processes in the occurrence/suppression of framing effects (e.g., Perez et al., 2018; Pu et al., 2017; Simon et al., 2004; Whitney et al., 2008). Three experimental approaches have been used to investigate this question. A first approach consists in investigating how individual differences in cognitive performances impact the sensibility to framing effects. Overall, individuals with higher cognitive performances on reasoning, inhibition, verbal intelligence, mathematics and memory were overall found to be less sensitive to framing manipulations (e.g., Bruine de Bruin, et al., 2007; 2012; Cokely, & Kelley, 2009; Kaczmarek et al., 2018; Perez et al., 2018; Smith & Levin, 1996). A second approach consists in promoting the extensive elaboration of the framed statements. Various studies have shown that when participants were encouraged to adopt an analytic reasoning to provide a rationale for their choices, framing effects were moderated or eliminated (Almashat, et al., 2008; Cheng et al., 2014; Kim, et al., 2005; Miller & Fagley, 1991; Sieck & Yates, 1997; Takemura, 1994; Thomas & Millar, 2012). A third approach consists in limiting participants’ ability to engage in effortful cognitive processing of the framed statement using either time pressure or cognitive load (e.g., Whitney et al., 2006). Various studies found that time pressure amplifies framing effects, as it affects the depth of the statement processing and favors a fast and intuitive decision process (Diederich & Wyszynski, 2017; Diederich et al., 2018; Guo et al., 2017; McElroy & Conrad, 2009). Some framing effect studies used dual-task paradigms to investigate the impact of cognitive load on the processing of framed statement (i.e., McElroy and Conrad, 2009; Whitney et al., 2008). Depleting participants’ cognitive resources using a competing task led to increased framing effects (McElroy & Conrad, 2009; Whitney et al., 2008). Taken together these results suggest that increasing responders’ cognitive load while playing the ultimatum game may lead them to reallocate cognitive resources from the affective reappraisal process to the processing of the problem statement, favoring the emergence of a framing effect.

Experiment 3 aimed at testing this assumption. In the study of Halali and colleagues (2014), participants had to perform a modified version of the Stroop task (1935) before playing the ultimatum game as responder. The modified Stroop task was used to increase responders’ cognitive load before they played the ultimatum game. As the present experiment was conducted on-line, we were not able to adopt such an experimental protocol due to its complexity. Instead, we chose to increase responders’ cognitive load by manipulating the complexity of the rules of the ultimatum game (e.g., Wendt, et al., 2014). Participants played a one-shot ultimatum game as in Experiment 2 but the statement of the economic game was made more complex using not one but two frame manipulations. The first one was an offer frame manipulation (similar to Experiments 1 and 2); and the second was an action frame manipulation where responders were asked which offers they would either accept or reject. We predicted that a greater cognitive effort would be necessary to process the double-framed problem compared to the simple-framed problem (Experiments 1 and 2) and may require responders to retrieve part of the cognitive resources previously allocated to the reappraisal of the negative affective response to the offers in the negative frame. This may affect the efficacy of the affective response reappraisal and allow the emergence of an offer framing effect, with responders being less prone to accept unfair offers in the offer negative frame than in the offer positive frame. We also predicted that we may observe an action framing effect with responders being more prone to accept unfair offers in the *accept* action frame (i.e., which offers would accept) than in the *reject* action frame (i.e., which offers would you reject). In other words, we predicted to observe a lower minimal acceptable offer 1) in the offer negative frame than in the offer positive frame; and 2) in the *accept* action frame than in the *reject* action frame.

### 4.1. Material and Methods

#### 4.1.1. Participants

436 French participants (194 females; mean age: 28 years ± 12) took part to this experiment. They received no compensation for their participation in the study.

#### 4.1.2. Ethics statement

All participants were informed of their rights, that they could withdraw from the experiment at any time and gave written informed consent for participation in the study, according to the Declaration of Helsinki. The research was carried out fulfilling ethical requirements in accordance with the standard procedures of the University of Modena and Reggio Emilia.

#### 4.1.3. Procedure

In Experiment 3, both the frames of the offer (i.e., negative frame: “*keep”* versus positive frame: “*offer*”) and the response to the offer (i.e., action frames: “*accept*” versus “*reject*”) were manipulated. The procedure was the same as Experiment 2 (first and second website pages). In the third website page, the ultimatum game statement was presented in the same way as Experiment 2, with the exception of the last sentence. For 114 participants (51 females), the final sentence of the problem was: “*If you were playing as responder, which amount(s) of money would you accept that the proposer keeps for himself ?”*, for the 103 participants (44 females) it was: “*If you were playing as responder, which amount(s) of money would you accept to be given by the proposer ?”*; for 111 participants (57 females), it was: “*If you were playing as responder, which amount(s) of money would you refuse that the proposer keeps for himself ?”*; for the 108 participants (42 females), it was: “*If you were playing as responder, which amount(s) of money would you refuse to be given by the proposer ?”*. The participants’ task was to select the icon(s) corresponding to the amount(s) of money they would accept versus refuse (i.e., 1€, 2€, 3€, 4€, 5€, 6€, 7€, 8€ or 9€) and to click on the icon “Terminate” to validate their choice after providing their age and biological gender and finally complete the experiment.

### 4.2. Results

The offers were analyzed in terms of minimal acceptable offer (i.e., the lowest offer the responders would accept). A 2 x 2 x 2 (Offer Frame [positive frame, negative frame] x Action Frame [“Accept”, “Refuse”] x Participants’ Gender [male, female]) ordinal logistic regression was performed on the minimal acceptable offer (1€, 2€, 3€, 4€, 5€, 6€, 7€, 8€, 9€). A manual stepwise analysis was performed to remove non-significant interactions from the model. The ordinal logistic regression analysis revealed a significant Offer Frame x Participants’ Gender [*B* (SE) = .716 (.352), CI (95%) = (.026, 1.406), Wald χ*²* (1) = 4.137, *p* < .05; see Figure 4] interaction with the offer positive frame predicting for a higher minimal acceptable offer (*M* = 4.04€) than the offer negative frame (*M* = 3.48€) in female participants but not in male participants (negative frame: *M* = 3.69€; positive frame: *M* = 3.75€).

**Figure 4.**
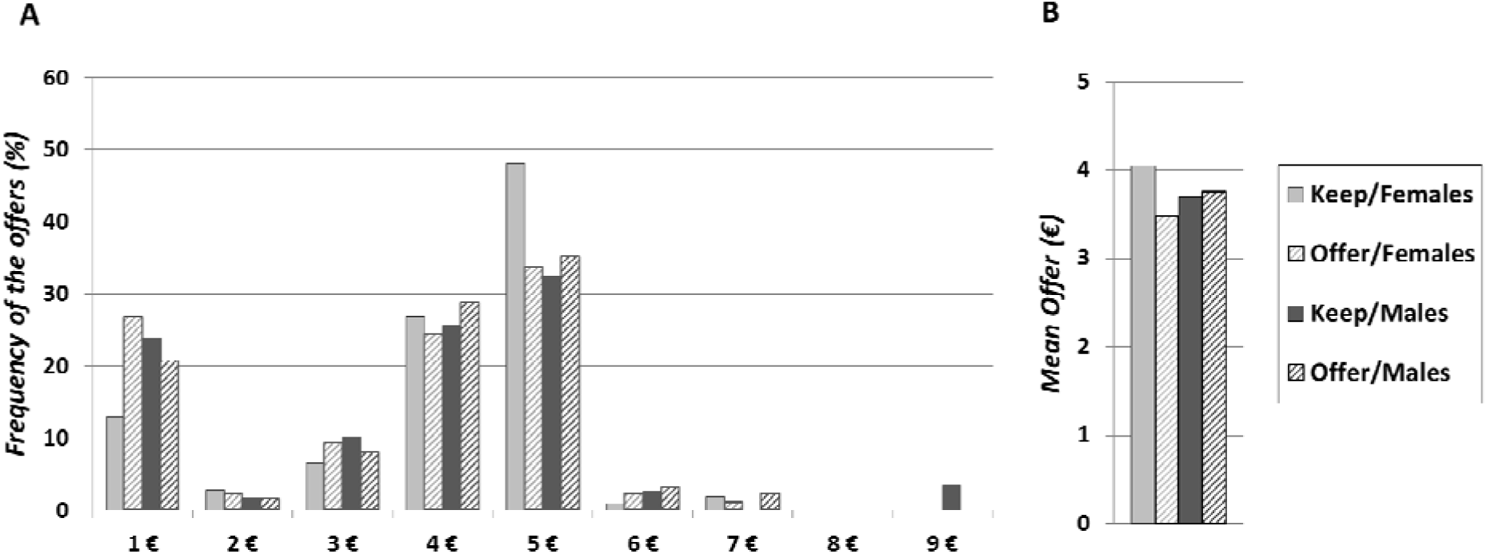
Illustrations of ***A)*** the frequency of minimal acceptable offers and the ***B)*** mean minimal acceptable offer in negative keep frame (plain) and positive offer frame (striped) for both female (light grey) and male (dark grey) responders.

The main effects of action frame [*B* (SE) = - .330 (.174), CI _(95%)_ = (- .671, .011), Wald χ*²* (1) = 3.588, *p* = .058], participants’ gender [*B* (SE) = - .250 (.254), CI _(95%)_ = (- .748, .248), Wald χ*²* (1) = .967, *p* = .325] and offer frame [*B* (SE) = - .129 (.235), CI _(95%)_ = (- .588, .331), Wald χ*²* (1) = .302, *p* = .583] did not reach significance.

### 4.3. Discussion

Experiment 3 was designed to test the hypothesis that increasing the complexity of the ultimatum game rules (i.e., double frame manipulation to increase cognitive load) may trigger a framing effect in responders. In order to increase the complexity of the ultimatum game, both the frame of the offer and the frame of the response (i.e., action frame) were manipulated. The results revealed an effect of the offer frame manipulation in female participants, with the negative frame predicting a higher minimal acceptable offer than the positive frame.

A first explanation for this result could be that processing a complex problem (i.e., double frame manipulation) was cognitively costly for the participants and that women were more sensitive to the offer frame manipulation because they have on average lower cognitive and computational capacities compared to men (e.g., Bruine de Bruin, et al., 2007, 2012; Cokely, & Kelley, 2009; Kaczmarek et al., 2018; Perez et al., 2018). This explanation may be very unlikely given that women were found to have comparable mathematical/computational performances as men when controlling for anxiety (Devine, et al., 2012) and that the task may not have triggered particular anxiety in participants (i.e., no pressure from the experimenter, no evaluation, anonymity of the response, possibility to quit the experiment).

Many studies found that women show greater emotional response to negative/aversive stimuli compared to men (e.g., Bradley, et al., 2001; Grossman & Wood, 1993; Strang et al., 2016). Therefore, another explanation could be that female responders showed a greater negative affective response in the negative frame compared to male responders. As the double-frame manipulation may have made the processing of the ultimatum game statement cognitively more demanding, the remaining cognitive resources may not have been sufficient to enable female responders to effectively reappraise the negative affective response in the negative frame due to its greater magnitude (i.e., respectively to male responders; Halali et al., 2013; Miu & Crişan, 2011), which resulted in a significant increase in female responders’ minimal acceptable offer in the negative frame compared to the positive frame. Overall, the results of economic game studies have shown that women tend to be more sensitive to both context modulations (Cox & Deck, 2006; Miller, & Ubeda, 2012) and unfairness than men (Croson & Gneezy, 2009; Fabre et al., 2015). Moreover, some framing effect studies reported gender differences regarding the sensitivity the framing manipulations, with women being more sensitive to frame manipulations than men in various life-and-death decision studies (e.g., Fagley & Miller, 1990, 1997; Fagley et al., 2010; Huang & Wang, 2010; Wang et al., 2001) and one mating decision study (Saad, & Gill, 2014). At the light of these previous studies, the results of the present experiment are not surprising.

In their repeated-measure study, Sarlo and colleagues (2013) found the opposite pattern of results, with male responders accepting less frequently mid-value offers in the negative frame than in the positive frame; and no effect of the framing manipulation on female responders. This difference in results could be due to a difference in the wording of the frame manipulations used in the present study and the study of Sarlo and colleagues (2013). While in the present study the negative frame “*I keep for myself*” and the positive frame “*I offer you*” were used, Sarlo and colleagues (2013) used “*I take*” as a negative frame and “*I give*” as a positive frame. The “*I take*” negative frame is quite particular as it depicts an active and quite aggressive monopolization of resources which could have triggered a strong negative reaction in males and inhibit women’s reaction, possibly explaining why they found a framing effect in males but not in females (Zeichner et al., 2003).

## 5. GENERAL DISCUSSION

Two previous studies investigated the impact of offer frame manipulations on the responders’ decision behavior, finding contradictory results (Sarlo et al., 2013; Tomasino et al., 2013). In the present study, three experiments were designed to further investigate this question. Experiment 1 used a similar experimental procedure as both the studies of Sarlo and colleagues (2013) and Tomasino and colleagues (2013). Participants played a repeated one-shot ultimatum game where they were proposed unfair, mid-value and fair offers framed either in a negative way (i.e., negative frame focusing on the money removed from the responder) or in a positive way (i.e., positive frame focusing on the money retrieved by the responder). In line with the study of Tomasino and colleagues (2013), no effect of the offer frame manipulations was found on responders’ acceptance rates in Experiment 1. However, longer response times were found in the negative frame than in the positive frame, reflecting a greater cognitive effort to decide on negatively framed offers than on positively framed offers. At a brain electrophysiological level, greater N400 amplitudes were found in response to the offers in the negative than in the positive frame, reflecting a greater negative affective response to the negatively framed offers (Bayer et al., 2010). This result confirms the findings of the two previous ultimatum game studies investigating framing effect that also found greater affective responses to the offers in the negative frame at both an electrophysiological level (Sarlo et al., 2013) and a neural level (Tomasino et al., 2013). Experiment 1 also showed greater P600 responses to 1) unfair offers than to both mid-value and fair offers; and 2) unfair offers in the negative frame than in the positive frame, suggesting that participants intensively reprocessed unfair offers, especially in the negative frame. This cognitive reprocessing appeared to have enabled the reappraisal of the negative affective response to the offers in the negative frame (Grecucci et al., 2013; Van’t Wout et al., 2010), which may have prevented the emergence of a framing effect (Miu & Crişan, 2011).

Experiment 2 was conducted to rule out the possibility that no framing effect was found at a behavioral level in both Experiment 1 and the study of Tomasino and colleagues (2013) because of their repeated-measures design. We made the assumption that repeated measurements could have triggered a habituation effect, reflected by a decrease in affective response over time, thus eliminating a potential framing effect. The experimental design of Experiment 2 was identical to the one of Experiment 1 at the exception that participants had only one economic interaction (i.e., unrepeated one-shot ultimatum game). Responders were found to be more likely to reject unfair offers and accept fair offers when playing the one-shot ultimatum game in its repeated form (Experiment 1; AR: 1€: 50.78 %; 3€: 61.20 %; 5€: 96.88 %) than in its unrepeated form (Experiment 2; AR: 1€: 6.83 %; 3€: 31.21%; 5€: 86.19 %). However Experiment 2 also failed to reveal an effect of the frame manipulation effect, suggesting that repeated measurements had no significant incidence on the affective response to framed offers in Experiment 1.

Experiment 3 aimed at investigating whether the failure to reveal a framing effect in Experiments 1 and 2 of the present study and the study of Tomasino and colleagues (2013) could be attributed to the relative simplicity of the ultimatum game problem. In order to affect the reappraisal of the negative affective response to negatively-framed offers and potentially trigger the emergence of a framing effect in the ultimatum game, the problem statement was made more complex to increase the responders’ cognitive load, using a double-frame manipulation consisting of an offer frame manipulation (similar to Experiments 1 and 2) and an action frame manipulation. The results showed a significant framing effect in female responders who asked for a higher minimal acceptable in the negative frame than in the positive frame. No framing effect was found in male participants. As women tend to show greater emotional response to negative/aversive stimuli compared to men (e.g., Bradley, et al., 2001; Grossman & Wood, 1993; Strang et al., 2016), the reappraisal of the negative affective response to negatively-framed offers may require greater cognitive resources for female than for male responders. This may be the reason why making the ultimatum game statement more complex (i.e., increased cognitive load to process the problem) may have affected female responders but not male responders.

Taken together these results suggest that the alternatives of a same problem may be associated with different cognitive costs and affective values, supporting the cognitive–affective tradeoff model (Gonzalez et al., 2005; Kahneman & Frederick, 2007). In the ultimatum game, offer frame manipulations − i.e., presenting the offers as the amount of money removed from versus received by the responder − appear to trigger different affective responses to the offers with more negative responses to the offers in the negative frame than in the positive frame (i.e., experiment 1; Sarlo et al., 2013; Tomasino et al., 2013), leading the responders to engage in a cognitively costly process aiming at reappraising the negative affective response to negatively framed offers (especially unfair ones). In the case the reappraisal process is effective, no framing effect may be observed (Experiments 1 and 2 of the present study; Tomasino et al., 2013). However, an offer framing effect might emerge (Experiment 3 of the present study; Sarlo et al., 2013) when responders are not able to allocate enough cognitive resources to the reappraisal process – either because these cognitive resources are allocated to a competing task or because they have low cognitive control abilities (i.e., “the ability to deliberately inhibit dominant, automatic, or prepotent responses to maximize the long-term best interests of an individual”; Halali et al., 2013).

Strong framing effects are usually found for decision problems with a high emotional resonance (for an overview, see Costa et al., 2014). Compared to a life-and-death problem such as the Asian disease problem (Tversky, & Kahneman, 1981), the ultimatum game has a low emotional resonance which allows the affective response in the negative frame to be reappraised more easily in the ultimatum game than in the Asian disease problem, explaining why most experiments failed to find an offer framing effect in the ultimatum game.

Similar to the previous ultimatum game studies investigating framing effect (Sarlo et al., 2013; Tomasino et al., 2013), the positive and the negative frames consisted in highlighting the money respectively received by the responder and removed from the latter. However, the wording of the frame manipulations used in the present study was not identical to previous studies. While in the present study the negative offer frame “*I keep for myself*” and the positive offer frame “*I offer you*” were used, the two previous studies used “*I take*” as a negative frame and “*I give*” as a positive frame. Experiment 3 showed that women are more sensitive to framing manipulations than men. However, Sarlo and colleagues (2013) using the “*I take*” / “*I give*” frames found that male responders (but not female responders) were sensitive to the offer frame manipulation. Further research is now needed to determine 1) whether different positive (e.g., “*I offer/give/leave you*”) and negative frames (e.g., “*I keep/take*”) modulate the strength of the framing effect in the ultimatum game and 2) whether men and women are diversely affected by these offer framing manipulations.

In the present study, Italian subjects were tested in Experiments 1 and 2 – as in the two previous studies using offer frame manipulations (Sarlo et al., 2013; Tomasino et al., 2013) – while the participants of Experiment 3 were French. While French and Italian are Latin languages with extremely close grammatical structures and vocabulary, we cannot be certain that cross-cultural differences do not trigger a variability in sensitivity to framing manipulations in the ultimatum game. Once again, further research is necessary to assess this potential limitation.

To sum up, the present study aimed at investigating the effects of offer frame manipulations on the decision behavior − and the associated electrophysiological brain correlates (Experiment 1) – of responders playing the ultimatum game. The results of Experiments 1, 2 and 3 suggest that the offer framing effect results from complex balance between affective and cognitive processes (in line with the cognitive–affective tradeoff model, Gonzalez et al., 2005; Kahneman & Frederick, 2007) and that the offer frame manipulations may not always lead to a framing effect due to the relatively low emotional resonance of the ultimatum game (Costa et al., 2014).

